# Burrow architectural types of the Atlantic ghost crab, *Ocypode quadrata* (Fabricius, 1787) (Brachyura: Ocypodidae), in Brazil

**DOI:** 10.1101/006098

**Authors:** Willian T. A. F. Silva, Tereza C. S. Calado

## Abstract

A broad range of aspects from paleontology to physiology of the ghost crabs *Ocypode quadrata* have been studied worldwide. These crabs have been used as ecological indicators of the levels of anthropogenic impacts on sandy beaches. Our aim is to report the variety of burrow architecture types constructed by ghost crabs *Ocypode quadrata* on beaches of Maceió, Brazil. We found 20 types of burrows that differ in shape (number of axes, number of openings, orientation of blind end, number of branches). The slash-shaped burrows (type C) were the most frequent shape, followed by types K (spiral) and E (Y-shaped). Type C also showed the largest opening diameter and length ranges. Burrow types F, J, P, S and T were the least frequent. The G-test for goodness of fit to a time-independent uniform frequency distribution (G = 417.61; d.f. = 18; p < 0.005) reject the hypothesis that burrow types are constructed randomly (uniform distribution). The dominance of type C burrows and other simple-type burrows over more elaborate types indicates preference for simplicity.

## INTRODUCTION

Ghost crabs are among the most terrestrial of the malacostracan crustaceans. They are represented by the genus *Ocypode* Weber, 1795 and are distributed worldwide in tropical and subtropical sandy beaches (Mclachlan and Brown, 2006). Their resistance to such a highly dynamic and variable habitat (affected by the high variation of temperature throughout the day and the continuous mechanical action of the waves) and their predatory dominance in the macro-infauna of these environments make them interesting organisms for studying the dynamics of sandy beaches.

Direct studies on ghost crab species have not been very common in the last few years due mainly to technical difficulties and low applicability. Most studies have been indirect, focusing on their burrowing behaviour instead of their population dynamics or organismal properties. However, a broad range of aspects, including paleontology (Curran and White, 1991; Portell et al., 2003), ecology (Fales, 1976; Loegering et al., 1995), biomechanics (Weinstein, 1995) and ecophysiology (Green, 1964; De Vries et al., 1994), of these organisms have been studied.

Spatial distribution, abundance and opening diameter of ghost crab burrows have been used as indirect estimators of population structure, age distribution and population size of these crabs on sandy beaches (Alberto and Fontoura, 1999; Araujo et al., 2008). Several studies have used ghost crabs burrows as bioindicators of levels of anthropogenic disturbances on sandy beaches and reported that the abundance of burrows is negatively correlated to the levels of anthropogenic impacts on the beaches under analyses (Warren, 1990; Peterson et al., 2000; Barros, 2001; Blankensteyn, 2006; Hobbs et al., 2008; Rosa and Borzone, 2008; Lucrezi et al., 2009). However, we recently showed that the number of ghost crab burrows might not represent the population size and must be interpreted with attention (Silva and Calado, 2013). And, despite the difference in abundance in urban and non-urban beaches, ghost crab burrows seem to show the same spatial distribution pattern (random distribution) (Silva and Calado, 2011).

Although anthropogenic impacts are an environmental issue, the ecological consequences of human impacts on sandy beaches, in particular the presence of off-road vehicles, need to be analyzed socio-culturally and economically in order to achieve an effective lasting system of conservation (Schlacher et al., 2007).

The burrowing behaviour shown by crabs is of extreme importance for their protection against predators and desiccation. Several studies have shown that burrow shapes vary from individual to individual and across species. Although it has been shown that bioturbation (the process of biological modification of sediments) has important implications for the biogeochemistry of the soil, soil biodiversity and ecosystem functioning (Meysman *et al*., 2006), studies on the architectural aspects of bioturbation (burrowing, more specifically) have been missed out.

Our main goal is to analyze the great variety of architectural types of burrows constructed by the Atlantic ghost crab *Ocypode quadrata* (Fabricius, 1787) on sandy beaches in Brazil and draw hypotheses to explain the variety and complexity of shapes shown. The burrowing behavior is an important part of crustacean biology that help us understand how these organisms interact with the environmental factors that compose their habitat.

## MATERIALS AND METHODS

This study was conducted between October 2006 and March 2007 non-periodically in two beaches, Avenida beach (9°40’16.8”S 35°44’36.4”W) and Pontal sand bar (9°42’08.4”S 35°46’56.9”W), in northeastern Brazil (Fig. 1). A total of 370 ghost crab burrows were excavated. This was accomplished using a trenching shovel and hand trowels. A long branch of *Ipomoea pes-caprae* (Convolvulaceae), popularly known as bayhops and morning-glory, was inserted into the burrow opening until resistance was met in order to determine tunnel direction and indicate the burrow floor in the event of a cave-in during excavation. Contrary to the method using plaster of Paris to analyze burrow architecture *in situ*, this method does not cause deformation of the fragile structure of the burrows. Since the researcher performs and visualizes directly the entire excavation process, deformation of burrow shape is controlled. Burrow length and opening diameter were measured with a 0.5 cm precision ruler. Sketches of burrow shapes were made for posterior digital enhancement. All burrows were backfilled upon the completion of the project. All burrows selected for excavation were located in a delimitated area of 600 square meters (6m in width and 100m in length) on the upper beach parallel to the shore line.

**Fig. 1.**
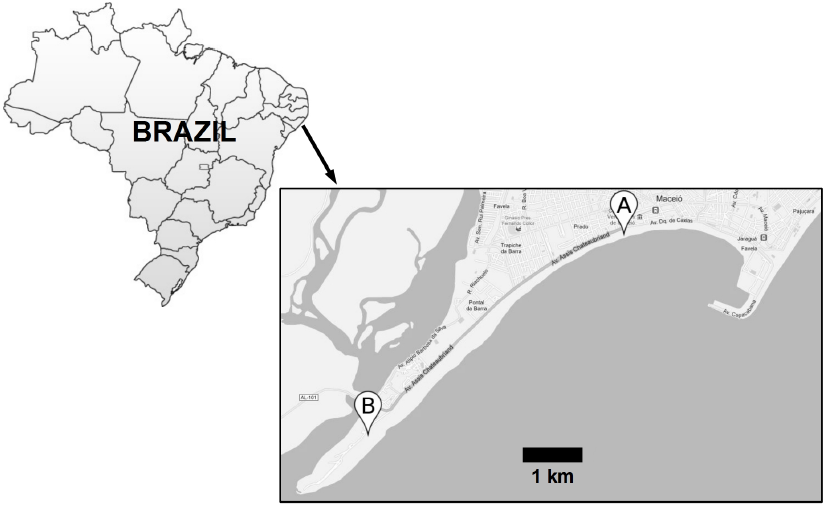
Localization of the beaches under study. A: Avenida beach. B: Pontal sand bar.

The data collected for burrows were used to construct graphs and calculate the measures of central tendency e variability. Due to the non-periodical sampling, the data obtained throughout the study period were pooled into a larger dataset, for statistical purposes. The G-test for goodness-of-fit is a likelihood-ratio test used to compare the fit of two models (the null model and the alternative model). It was computed to test the null hypothesis that the observed frequencies of burrow types over the study period follow a time-independent uniform frequency distribution (null model). The Pearson product-moment correlation coefficient was calculated as an approximation of the population parameter ρ to measure the strength of association between burrow length and burrow opening diameter. It was assumed that larger opening diameters would be associated with longer lengths, then burrow opening diameter was designated as the X variable and burrow length was designated as the Y variable. The null hypothesis (ρ=0) states that there is no association between burrow length and burrow opening diameter. The alternative hypothesis (ρ>0) states that there is a positive association between the two variables.

## RESULTS

Our excavations revealed that ghost crab burrows vary in shape from simple slash-shaped burrows (type C) to relatively complex branched and multiple-opening shapes (e.g. type I). Twenty types of burrow architecture were identified (Fig. 2), including J-shaped, Y-shaped, U-shaped, V-shaped, spiral and single tube burrow types. A complexity level, from I to III, was assigned to each burrow type: level I was assigned to all burrow types with a single opening and no branches; level II was assigned to types with two openings and no branches, and to types with a single opening and bearing branches; level III was assigned to types with two openings and bearing branches. The great majority of burrow types (84.32%) fall within complexity level I (Fig. 3). The slash-shaped burrows (type C) were the most frequent shape (N = 211), followed by types K (spiral) and E (Y-shaped). The G-test for goodness of fit to a time-independent uniform frequency distribution (G = 417.61; d.f. = 18; p < 0.05) reject the hypothesis that burrow types are uniformly distributed. Type C burrows also showed the largest opening diameter and length ranges (Fig. 4), having a mean diameter ± SD of 2.05 ± 0.76 cm and a mean length ± SD of 51.59 ± 25.64 cm. Type C burrows consist of a single straight tube inclined down from the surface. Burrow types F, J, P, S and T were the least frequent, being found only one unit of each of these types, and for which means and standard deviations were not calculated. Type A burrows had the largest mean diameter of 2.35 ± 0.47 cm, type G having the greatest mean length of 74 ± 11.31 cm. Variances of burrow length and diameter were 625.23 and 0.52, respectively, and the coefficients of variation for burrow length and diameter were 45.64% and 35.54%, respectively, for the entire study period. The Pearson product-moment correlation coefficient (r = 0.28; one-tailed p < 0.05) support the alternative hypothesis (ρ>0) that there is a significant positive association between burrow length and burrow opening diameter. However, the coefficient of determination (r^2^ = 0.08) is very low, indicating a very weak linear association between the two variables (Fig. 5)

**Fig. 2.**
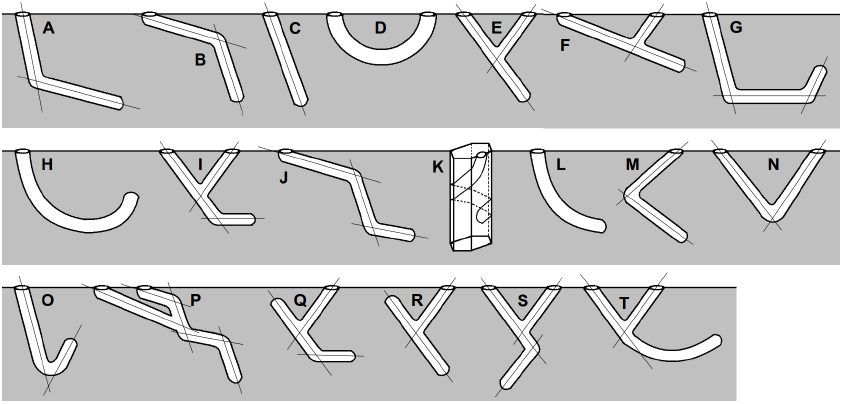
Architectural types of ghost crab burrows. Straight lines represent the axes upon which the burrow architecture is constructed. Letters were randomly assigned to burrow types to facilitate data assortment.

**Fig. 3.**
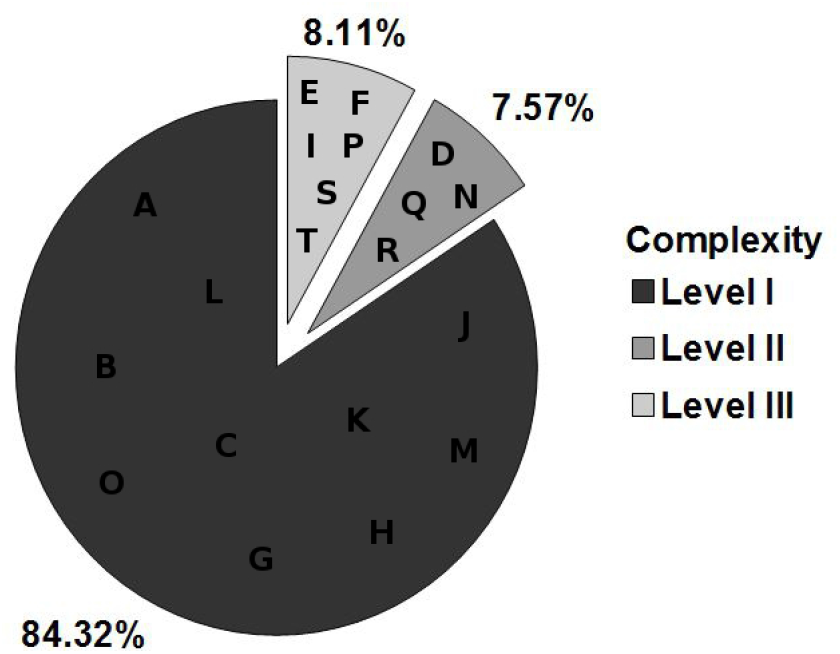
Frequency of burrow type complexity levels, according to the classification of complexity used in the present study.

**Fig. 4.**
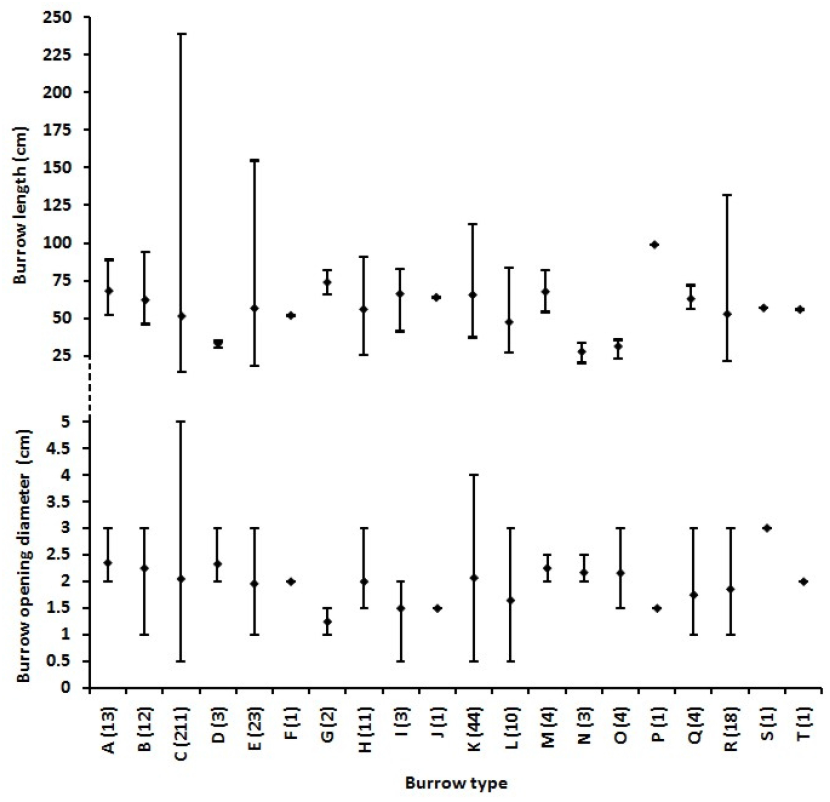
Mean opening diameter and length of ghost crab burrows. Vertical bars represent the minimum and maximum values. N is in parenthesis.

**Fig. 5.**
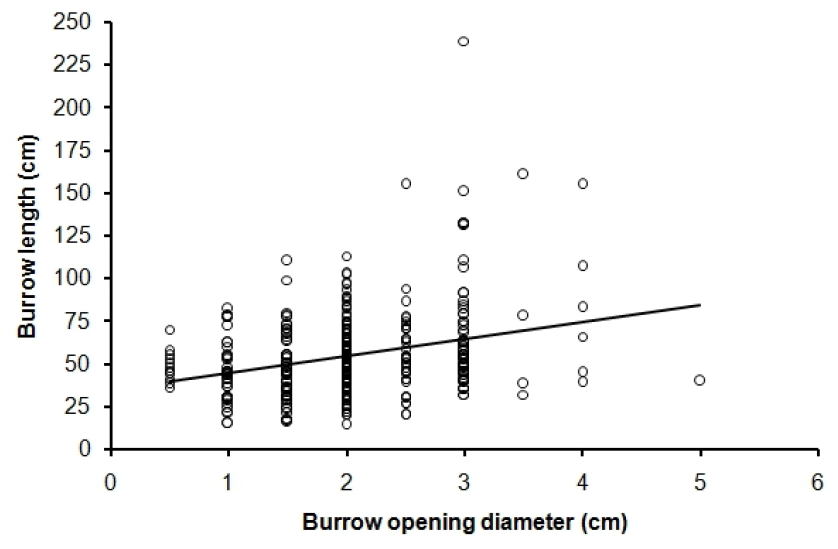
Scatter plot with regression line of burrow opening diameter and burrow length. Pearson product-moment correlation coefficient (r = 0.28; one-tailed p < 0.05) and coefficient of determination (r^2^ = 0.08).

Different burrow architecture types were defined according to the number of openings to the outside, the number and position of the axes (straight lines) that form the burrow shape, the existence of branches, and the orientation of the spherical blind end (Table 1). Seven (types D, E, F, I, N, P, S and T) out of the 20 types found have two aboveground entrances, the remaining types having only one entrance. Twelve (types A, B, C, D, G, H, J, K, L, M, N and O) out of the 20 types are single tube burrows, the remaining types being branched burrows. Number of axes varies from one to five, 2-axis structures making the majority of burrow types. All burrow types but D and N terminate in one or two spherical blind ends. Types D, H and L consist of different types of arched single tubes. Types E, F, P, Q, R, S and T consist of different morphologies of Y-shaped burrows. Types E and F differ only in the symmetry of the upper arms. In types Q and R, the upward branch does not extend up to the surface and terminates in a spherical blind end. Burrows that have the same number of openings, axes and branches, and same orientation of the blind ends (e.g. A, B and M; E and F) distinguish from one another by the arrangement of their axes.

**Table 1.** Burrow type characterization according to number of openings, number of axes, presence of branches, and orientation of the dead end (when present). A = arched axis; S = spiral axis.

| Burrow type | Number of openings | Number of axes | Branched? | Blind end |
| --- | --- | --- | --- | --- |
| A | 1 | 2 | No | Downward |
| B | 1 | 2 | No | Downward |
| C | 1 | 1 | No | Downward |
| D | 2 | A | No | Absent |
| E | 2 | 2 | Yes | Downward |
| F | 2 | 2 | Yes | Downward |
| G | 1 | 3 | No | Upward |
| H | 1 | A | No | Upward |
| I | 2 | 3 | Yes | Parallel to superficial plane |
| J | 1 | 3 | No | Downward |
| K | 1 | S | No | Downward |
| L | 1 | A | No | Downward |
| M | 1 | 2 | No | Downward |
| N | 2 | 2 | No | Absent |
| O | 1 | 2 | No | Upward |
| P | 2 | 5 | Yes | Downward |
| Q | 1 | 3 | Yes | (2) Upward; parallel to superficial plane |
| R | 1 | 2 | Yes | (2) Upward; downward |
| S | 2 | 3 | Yes | Downward |
| T | 2 | 2 + A | Yes | Upward |

## DISCUSSION

*Ocypode quadrata* construct a great variety of burrow architectural types. Several factors (for example age group, cephalothorax size, granule size, sand composition) may influence the complexity of the architectural types preferred by crabs and further studies need to be conducted in order to determine which factors are important. However, the higher frequency of simple type burrows over complex type burrows indicates preference for simplicity. Because burrowing is energetically costly both for invertebrates (e.g. Brown, 1979) and for vertebrates (e.g. Du Toit et al., 1985) and complexity is associated with higher energetic costs, preference for simplicity may represent an evolutionary strategy.

Additionally, in an evolutionary context, one could speculate whether burrow architectural types or the ability to construct deeper burrows are under selective pressure. This speculation arises from the frequency distribution of architectural types observed here. For example, type C burrows (and complexity type I burrows in general) represent over 50% of all burrows on a beach. With regard to burrow depth (length), deeper burrows could increase protection from potential predators and desiccation, which would explain selection upon burrow depth. However, given the association between burrow diameter (an estimate of crab size) and burrow length, it is safer to assume that burrow length is associated to ontogeny, rather than a behavioral trait that is subject to selection. Also, the high variance of vertical length of burrows is not a positive signal that indicates the action of selective pressure upon burrowing behavior.

According to Shuchman and Warburg (1978), burrow length of *O. cursor* varies with distance from the water, deeper burrows being located further away from the water than shallower burrows. Also, burrows are distributed by size and density along the beach width and inclination. Similarly, number and diameter of burrows of *O. quadrata* tend to increase as the distance from the water line increases (Duncan, 1986; Araujo et al., 2008). Since there is a strong relationship between the crab carapace width and its burrow opening diameter in *O. quadrata* (Alberto and Fontoura, 1999), as well as between the crab carapace length and its burrow length and opening diameter in *O. ceratophthalma* (Chan et al., 2006), the aforementioned studies indicate that younger crabs usually construct their burrows in the lower beach whereas older crabs construct their burrows in the upper beach, suggesting a relationship between distance from sea water and ontogeny. Ghost crab burrows are thus indicative of three age zones within the supra-littoral zone of sandy beaches (Frey, 1970).

When architectural types were compared across studies in different *Ocypode* species (*O. quadrata*, *O. cursor* and *O. ceratophthalma*), we found that, in addition to some of the types described in this study, other architectural types had been described (e.g. Shuchman and Warburg, 1978; Chakrabarti, 1981; Chan et al., 2006).

Burrows of *Ocypode ceratophthalma* are similar in shape and size to burrows of *Ocypode quadrata*. However, the former species shows more complex burrow architectural types, with secondary branching (Chakrabarti, 1981) and chambers (Chan et al., 2006). Chambers were not found at the base of the burrows in this study and no other study has reported the presence of such chambers. The chambers found by Chan et al. (2006) might have resulted from the accumulation of plaster of Paris used in their research methods.

Type R burrows (Y-shaped) are the most common type constructed by *O. cursor*, rather than type C burrows that represented the majority of *O. quadrata* burrows. Like in *O. quadrata*, *Ocypode cursor* type R burrows consist of a long arm that opens to the surface and a short arm that extends towards the surface but ends blindly. Type E burrows (Shuchman and Warburg, 1978) and J- and L-shaped burrows (Strachan et al., 1999) are also constructed by *O. cursor* but the former represent only a small percentage of all burrows.

According to Frey (1970), Y-shaped and U-shaped burrows are usually constructed by crabs of intermediate age, whereas single tube burrows oriented nearly vertically in the substrate are typical of young crabs. Burrows constructed by old crabs are larger but not usually branched.

Trace fossils of *O. quadrata*, denominated *Psilonichnus upsilon*, have been reported to show the same major shape types, e.g. J-shaped, U-shaped and Y-shaped types, found in the present study. Because of the strong relationship between these trace fossils and their trace-makers, they are useful as indicators of past sea-level position (Curran & White 1991).

The several types of burrows constructed by ghost crabs and several other species raise important questions that, if answered, would facilitate the understanding of the ecology of these species. For example, are burrow architectural types constructed at random? or is each type of burrow used for a specific reason? Some researchers suggest that ghost crabs construct different types of burrows that are used for different functions throughout their life span (Chan et al., 2006). According to Clayton (2005), spiral burrows are constructed by male *O. jousseaumei* crabs each tidal cycle and are used for courtship. However, in *O. ceratophthalma*, architectural types of ghost crab burrows do not differ between sexes, according to Chakrabarti’s (1981) study, but this study did not report any spiral burrow.

Clayton (2005) also found an association between the handedness of the crabs’ major chela and the direction of spiralization of their burrows, and Duncan (1986) found that ghost crab burrows have a non-random orientation, following the orientation of the beach slopes and dune faces. These findings support the hypothesis of non-random construction of different architectural types of burrows.

Barrass (1963) distinguished two types of burrows according to the presence or absence of a pile of sand at the opening of the burrows, indicating whether the burrow was used as an entrance (called true burrows) or exit (called emergency holes), respectively. As most burrow types found in the present study showed only one opening, these openings cannot be classified as entrance or exit holes.

Ghost crab burrows also show differential patterns of distribution according to their age group. Adult ghost crab burrows are uniformly distributed, whereas juvenile crab burrows show an aggregated pattern of distribution. This difference in distribution between age groups may reflect the physiological competence of young and old crabs to resist desiccation (Fisher and Tevesz, 1979).

Understanding how ghost crabs construct their burrows and how each architectural type of burrow is used is very important to understanding the biology of these organisms. Since the *Ocypode* genus has been used as a bioindicator of the environmental quality, this type of information is not only useful for scientific reasons but also for the development of better management and conservation strategies of sandy beaches, through a better understanding of the biology of these bioindicators and their interaction with the environment. However, probably due to the difficulty of studying such organisms, they have not been extensively studied.

## Acknowledgements

We are grateful to all members of LabMar at Federal University of Alagoas for enthusiastic support. Special thanks are due to Manoel J. Silva, José Jonathas P. R. Lira, Marianna B. Baptista and Liliane S. S. Tonial for help with the fieldwork and to Susanne Zajitschek for her critical review of the manuscript and suggestions for its improvement.

